# Male density and rapid evolution of genital morphology in the seed beetle *Callosobruchus maculatus*

**DOI:** 10.1101/052332

**Authors:** D. M. Soper, W. L. Macy

## Abstract

Male reproductive structures are known to be extremely diverse, particularly in insect taxa. Male genital structures are thought to be some of the fastest evolving traits, but the processes responsible for this pattern remain unclear. In the present study we manipulated the mating regimes of *Callosobruchus maculatus*, a seed beetle, to determine if male genital structures would be altered under forced monogamy and polyandry. Males in this species have an intromittent organ that contains spines that are known to puncture the female reproductive tract. We measured both testes size and genital spine length in monogamous and polyandrous treatments over seven generations. We found that testes size was not significantly different between treatments, but that genital spine length was significantly longer in the polyandrous treatment within seven generations. These results highlight the fact that evolution can occur rapidly when under strong sexual selection, a process that has been implicated in leading to morphological differences in male genitalia.

## Introduction

Male genitalia of internally fertilizing species are extremely diverse and thought to evolve at a rapid pace based on phylogenetic species comparisons [1]. In fact, insect male genitalia are so diverse compared to other morphological characters that taxonomists have used the structure for species identification and classification [2]. Several studies have attempted to determine the cause for such a plethora of diversity; current evidence appears to overwhelmingly support hypotheses under the umbrella of sexual selection [3, [4].

Mating system of a species can influence the strength of sexual selection [5]. For example, polyandry may strengthen sexual selection because males compete with one another at the cellular level through sperm competition [6], at the organismal level through direct combat [7], and/or at the organ level, whereby genital shape may influence sperm transfer and the number of fertilization events [2]. Although current evidence supports the idea that sexual selection is responsible for the widespread diversity of male genitalia and suggests that genital evolution can occur rapidly [4], direct observation of morphological evolution remains scant.

Here we track the evolution of male genitalia in *Callosobruchus maculatus*. In this sexually dimorphic seed beetle, males have testes and a seminal vesicle that leads to an intromittent organ with sclerotized spines (Fig. 1). Spines are known to puncture the reproductive tract of the female, possibly as a way to inject antiaphrodisiacs and shorten female life span [8]. Using experimental evolution, we measured male genital spines, testicle size, and body size over seven generations under two mating regimes of monogamy and polyandry. We predicted that increased spine length and testes size would occur in the polyandry treatments due to increased sperm competition.

**Figure 1.**
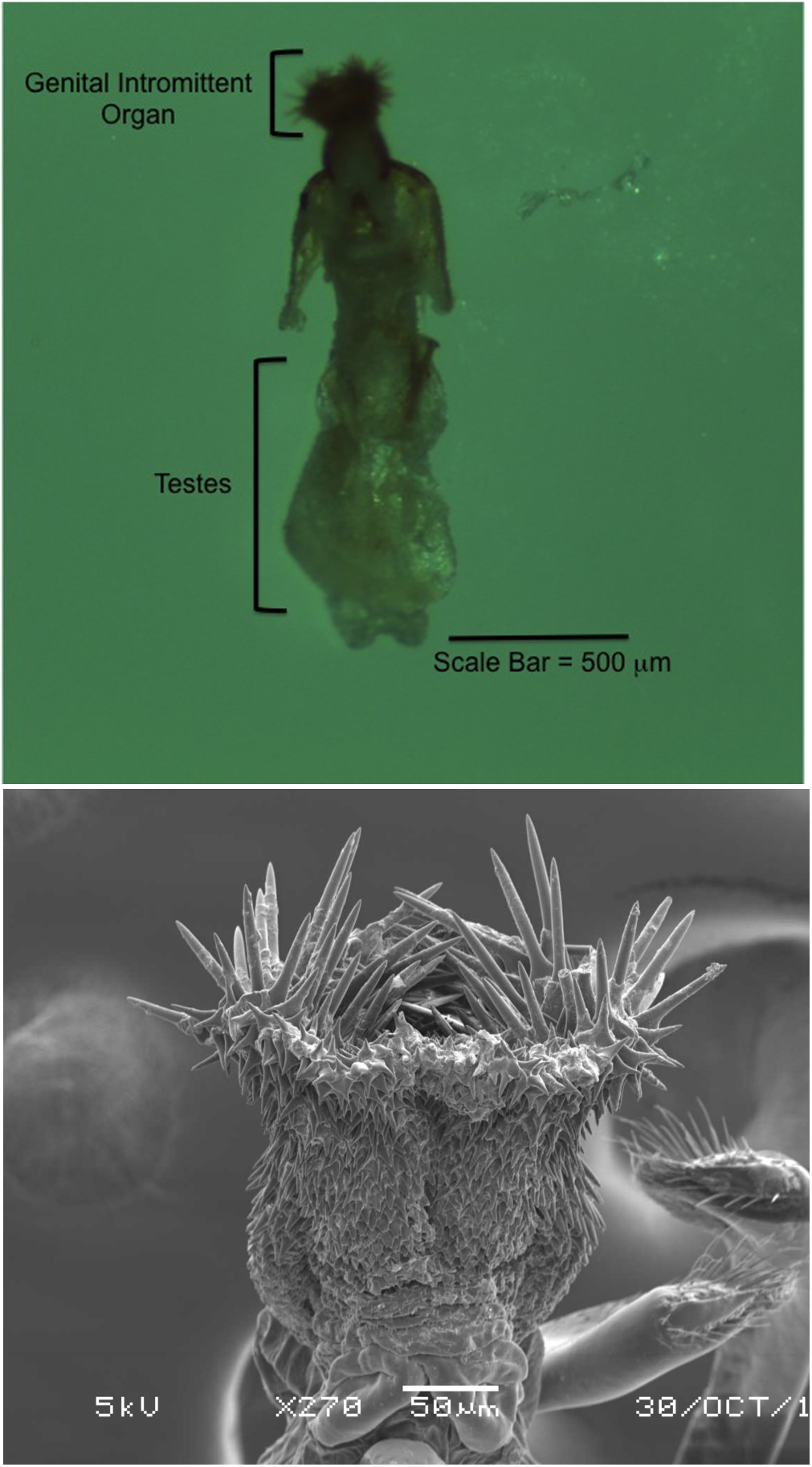
The anatomy of the *Callosobruchus maculatus* male reproductive tract. (*a*) Testicular structure shown with male intromittent organ in the everted state exposing spines under a dissecting microscope *(b)* Male intromittent organ tip showing spines under scanning electron microscope. Scale bar, 500mm in *(a)*, and 50mm in *(b)*.

## Materials and Methodss

### Study System

*Callosobruchus maculatus*, a cosmopolitan pest, was used as the study system because they are sexually dimorphic, easy to care for, and have a generation time of 3-4 weeks. Eggs are laid on bean hosts and after 4-8 days the larvae hatch and burrow into the beans where the beetles develop into adults. Adult beetles live approximately two weeks without the need for food or water.

### Treatments

This experiment used two levels of male-male mate competition as treatments. Monandrous treatments had no male competition because they consisted of one male and one female. Polyandrous treatments had intense male competition because they consisted of six males and one female. Carolina Biological (Burlington, NC) supplied the initial generation of beetles. Mating groups were created by randomly assigning individuals to a mating pairings within its treatment, and half-sib inbreeding was avoided.

Mating pairs were placed in 35mm Petri dishes, and were observed for 20 minutes before dried mung beans were added for the females to use for egg laying. This observation period allowed for behavioral data collection that will be published independently. Beetles remained in their dishes for 7 to 8 days before being removed. The beans with eggs were isolated before emergence to yield virgin beetles for the next generation. Once the new beetles emerged (approximately every 4 weeks), they matured 48 h before being placed into mating groups. This procedure was repeated for a total of seven generations of offspring from the initial populations.

### Data Collection

Starting at generation three, 12 males from each generation were randomly selected from each treatment for data collection. Body size, testicular size, and average intromittent organ spine length was collected from each individual. Dissection, body images, and testicle images were all generated with a Leica EZ4HD dissecting microscope. Intromittent organ spine images were captured with a JEOL JSM-5900LV scanning electron microscope, and images were then analyzed using Image J v. 1.48 [9].

#### Body and Testicle Size

Beetles were placed on ice for approximately 1 minute to anesthetize the organisms to allow for microscopy and dissection. If the beetle regained activity prior to dissection, the organism was placed back on ice for an additional minute. Body size was measured from images taken prior to dissection and calculated as the area created by multiplying the width of the elytra by the length of the elytra at their widest points. Dissection for removal of the reproductive organs occurred by submerging the beetle in 50μl of buffer solution that consisted of 1.4 M NaCl, 0.03M KCl, 0.1 M Na2HPO4, and 0.02 M KH2PO4 (Sigma-Aldrich, St. Louis, MO). Testicle images (fig. 1A) captured after the removal of the seminal vesicle were measured to obtain a testicle area for each male.

#### Intromittent Organ Spine Length

Prior to imaging, gentle pressure was applied to the base of the intromittent organs so that spines, which are folded inward when stimulation is not present, were exposed. The organs were then dried and mounted for imaging. Images of the extended intromittent organ were taken on a scanning electron microscope for enhanced magnification. Mean individual spine length was calculated with 5-10 randomly selected tip spines from each individual.

## Statistical Analysis

Body size was first checked for correlation with both testicle size and intromittent organ spine length to determine whether larger beetles had larger testes or longer spines. After data was analyzed for normality, both testicle size and intromittent organ spine length were analyzed with an Analysis of Variance with interaction between generation and treatment. ANOVAs were used to determine if there was a significant difference in the means of our treatments across the seven generations. All statistical analysis was conducted with JMP Statistical Discovery, from SAS (Cary, NC).

## Results

Mating regime did not affect testicle size (F= 0.016, p = 0.9181). There was also no correlation of body size to testicle area (R^2^= 0.006, p = 0.955) or genital spine length (R^2^ = 0.069, p= 0.694). We did find evidence for an evolved relationship between mating regime and genital spine length: When averaged across generations, polyandrous males had significantly longer spines than monandrous males (Fig. 2A, Polyandrous mean = 1.2 μm, Monandrous mean= 1.0 μm, F = 4.27, p = 0.01). Both treatment (fig. 2B, F = 4.87, p = 0.03)and generations (Fig. 2B, F = 6.93, p = 0.01) significantly interacted with genital spine length, driven by the increase in polyandrous male spine length over the seven generations of the experiment. By generation seven, egg production declined in females in the polyandrous treatment to an extent that led to the termination of the experiment due to the lack of ability to maintain six males to one female without inbreeding. This phenomenon did not occur in the forced monogamy treatment.

**Figure 2.**
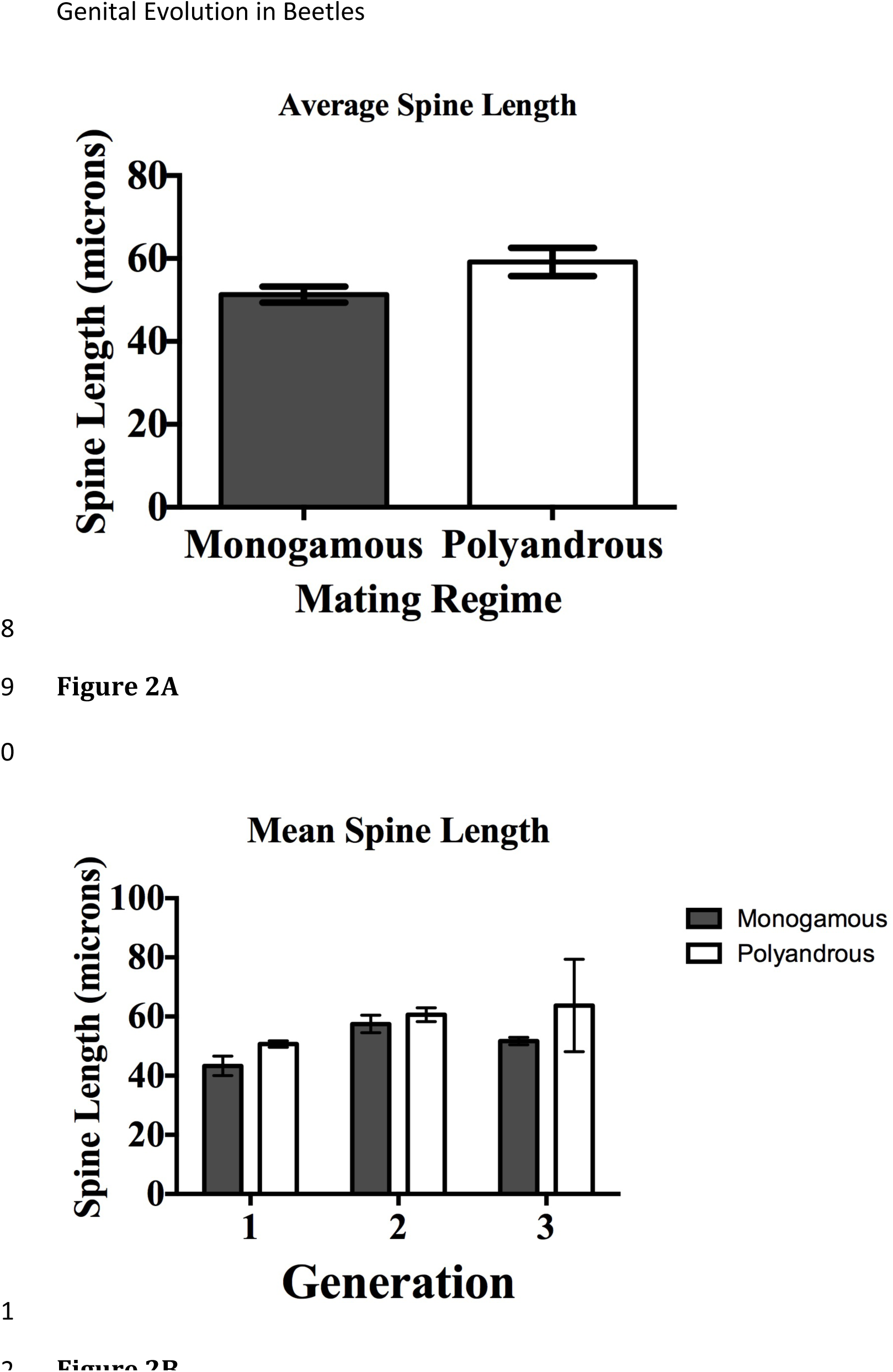
Intromittent spine length (*a*) the effect of treatment (monandrous treatment grey bars, polyandrous treatment white bars) on spine length and *(b)* the effect of generations and treatment on average spine length.

## Discussion

After seven generations, we found that males experiencing intense male competition had significantly longer sclerotized spines on genital intromission organs than males that experienced no competition from conspecifics. Body size did not correlate with either spine length or testes size in either mating regime. These results suggest that mating regime, which itself might be driven by population sex ratio, could influence the evolution of genital morphologies and influence the fitness of individuals within populations. The polyandrous treatment experienced a decline in the number of viable offspring until generation seven, when numbers were reduced to the point of forcing the termination of the study.

The genital spines of *Callosobruchus maculatus* are known to produce extensive genital damage in females, which can lead to reduction in fitness [10], but also are necessary to alter the female’ s physiology to increase the mating male’ s chance of fertilization [11]. *Callosobruchus maculatus* males with long spines are more successful at fertilization and inject significantly more seminal fluid into female hemolymph [11], which can positively influence fertilization success but also may lead to negative fitness effects particularly for females that mate multiply due to genital damage and reduced female longevity [8,[12]. It is possible that the increase in length observed for the polyandrous treatment may have contributed to extensive damage and reduction in female fecundity. Moreover, female life span may have been reduced leading to fewer eggs being produced.

A similar study also using *C. maculatus* found that forced monogamy resulted in shorter male genital spines, while a polygamous (multiple males and multiple females) male genital spine length was maintained [13]. Although the present study also had a forced monogamy treatment, here we found that spine length was maintained in monogamy and under polyandry (one female and multiple males) spine length increased. This difference between study outcomes may have occurred because Cayetano et al. (2011) was carried out to 18-21 generations, while the present study was performed to generation 7. Therefore, it is possible the same result for the monogamous treatment would have been achieved if the study were continued for more generations. Spine length increase may not have been observed under polygamous mating because males had the opportunity to mate with multiple females. Whereas in the present study, under polyandry, a male was required to outcompete other males in order to be successful at fertilization. This increase in sexual selection pressure could have been responsible for the increase in spine length.

Another reproductive trait that is commonly positively correlated with increased male competition is testicle size [14,[15] because typically larger testes results in higher sperm counts. Counter to this expectation, we found that testicle size did not differ between monogamous and polyandrous treatments. This may be due to the fact that *C. maculatus* males produce and inseminate more sperm than can be possibly stored in the female spermathecae [16], and thus, *C. maculatus* males already produce the maximum amount of sperm that is possible regardless of the level of male competition that is present. Excess sperm deposition in this species is thought to occur because large numbers of sperm deposited in the female spermathecae increases the time with which she will re-mate, which allows more time for fertilization to occur [16].

Male reproductive morphologies are known to be some of the fastest evolving characters due to their direct effect on fitness [1]. In insects, this has led to a wide diversity of genital morphologies including a breathtaking array of intromittent organs [17] that have even been used for species identification. From lock-and-key mechanisms in damselflies [18] to twin claw-like genital structures in *Drosophila* [19], genital intromittent organs occur in a variety of structures for seemingly various reasons. Sexual selection has been the primary force suggested as to how these structures can evolve such diversity, but observation of rapid genital change has been rare. Lack of direct evidence that male competition can lead to genital morphological alterations means that if and/or how this process occurs remains obscure. The present study shows that genital intromittent organ evolution can occur rapidly, under intense male competition.

## Acknowledgements

We would like to thank the Beloit Biology Department, especially Y. Grossman for statistical support and guidance. We also would like to thank the Beloit College Chemistry department for assistance and training in use of the Scanning Electron Microscope.

## Data Accessibility

Data to support figure 2A & 2B will be deposited to Dryad upon acceptance.

## References

1. Arnqvist G. 1998 Comparative evidence for the evolution of genitalia by sexual selection. Nature393, 784–786.

2. Schilthuizen M. 2003 Shape matters: the evolution of insect genitalia. Proc. Exper. Appl. Entomol, NEVAmsterdam. 14, 9–16.

3. Hosken D.J., Stockley P. 2004 Sexual selection and genital evolution.Trends Ecol. Evol 19, 87–93.

4. Simmons L.W. 2014 Sexual selection and genital evolution. Austral Entomology 53, 1–17.

5. Kokko H., Rankin D.J. 2006 Lonely hearts or sex in the city? Density-dependent effects in mating systems.Phil. Trans. R. Soc.B 361, 319–334.

6. Birkhead T.R., Møller A.P. 1998 Sperm competition and sexual selection, Academic Press.

7. Moczek A.P., Emlen D.J. 2000 Male horn dimorphism in the scarab beetle, Onthophagus taurus: do alternative reproductive tactics favour alternative phenotypes? Anim. Behav. 59, 459–466.

8. Crudgington H.S., Siva-Jothy M.T. 2000 Genital damage, kicking and early death. Nature 407, 855–856.

9. Schneider C.A., Rasband W.S., Eliceiri K.W. 2012 NIH Image to ImageJ: 25 years of image analysis. Nature methods 9, 671–675.

10. Gay L., Hosken D.J., Eady P., Vasudev R., Tregenza T. 2011 The evolution of harm–effect of sexual conflicts and population size. Evolution 65, 725–737.

11. Hotzy C., Polak M., Rönn J.L., Arnqvist G. 2012 Phenotypic engineering unveils the function of genital morphology. Curr. Biol. 22, 2258–2261.

12. Savalli U., Fox C. 1999 The effect of male mating history on paternal investment, fecundity and female remating in the seed beetle Callosobruchus maculatus. Funct. Ecol. 13, 169–177.

13. Cayetano L., Maklakov A.A., Brooks R.C., Bonduriansky R. 2011 Evolution of male and female genitalia following release from sexual selection. Evolution 65, 2171–2183.

14. Gage M.J. 1994 Associations between body size, mating pattern, testis size and sperm lengths across butterflies. Proc. R. Soc. B. 258, 247–254.

15. Pitcher T.E., Neff B.D., Rodd F.H., Rowe L. 2003 Multiple mating and sequential mate choice in guppies: females trade up. Proc. R. Soc. B. 270,1623.

16. Eady P. 1995 Why do male Callosobruchus maculatus beetles inseminate so many sperm? Behav. Ecol and Socio. 36, 25–32.

17. Shuker D.M., Simmons L.W. 2014 The evolution of insect mating systems. New York, NY, Oxford University Press, USA.

18. McPeek M.A., Symes L.B., Zong D.M., McPeek C.L. 2011 Species recognition and patterns of population variation in the reproductive structures of a damselfly genus. Evolution 65, 419–428.

19. Kamimura Y. 2007 Twin intromittent organs of *Drosophila* for traumatic insemination. Biol. Lett. 3, 401–404.

